# *Dictyostelium discoideum-*assisted pharmacognosy of plant resources for discovering antivirulence molecules targeting *Klebsiella pneumoniae*

**DOI:** 10.1101/2023.10.27.564015

**Authors:** Marcos Hernández, Carlos Areche, Grover Castañeta, Diego Rojas, Macarena A. Varas, Andrés E. Marcoleta, Francisco P. Chávez

**Affiliations:** Laboratorio de Microbiología de Sistemas, Departamento de Biología, Facultad de Ciencias, Universidad de Chile, Santiago, Chile; Departamento de Química, Facultad de Ciencias, Universidad de Chile, Santiago, Chile; Grupo de Microbiología Integrativa, Laboratorio de Biología Estructural y Molecular, Departamento de Biología, Facultad de Ciencias, Universidad de Chile, Santiago, Chile

**Keywords:** . Antimicrobials, multidrug-resistant pathogens, antivirulence, plant extracts, pharmacognosy

## Abstract

The rise of antibiotic-resistant bacterial strains poses a significant global health challenge, underscoring the critical need for innovative strategies to address this threat. Natural products and their derivatives have emerged as a promising reservoir for drug discovery. The social amoeba *Dictyostelium discoideum* is an advantageous model organism in this effort. Using this invertebrate model, we introduce a novel perspective to screen natural plant extracts for molecules with potential antivirulence activity. As a proof of concept, we established a simple high-throughput assay to screen for antivirulence molecules targeting *Klebsiella pneumoniae* among extracts of *Helenium aromaticum*. Thus, we aimed to identify compounds attenuating *K. pneumoniae* virulence without inducing cytotoxic effects on amoeba cells. Notably, the methanolic root extract of *H. aromaticum* but not other extracts fulfilled these prerequisites. Further analysis via UHPLC-ESI-MS/MS led to the identification of 24 chemical compounds boasting potential antivirulence attributes. This research underscores the potential of employing *D. discoideum*-assisted pharmacognosy for unearthing novel antivirulence agents against multidrug-resistant pathogens.

**Table of Content Graphic:** 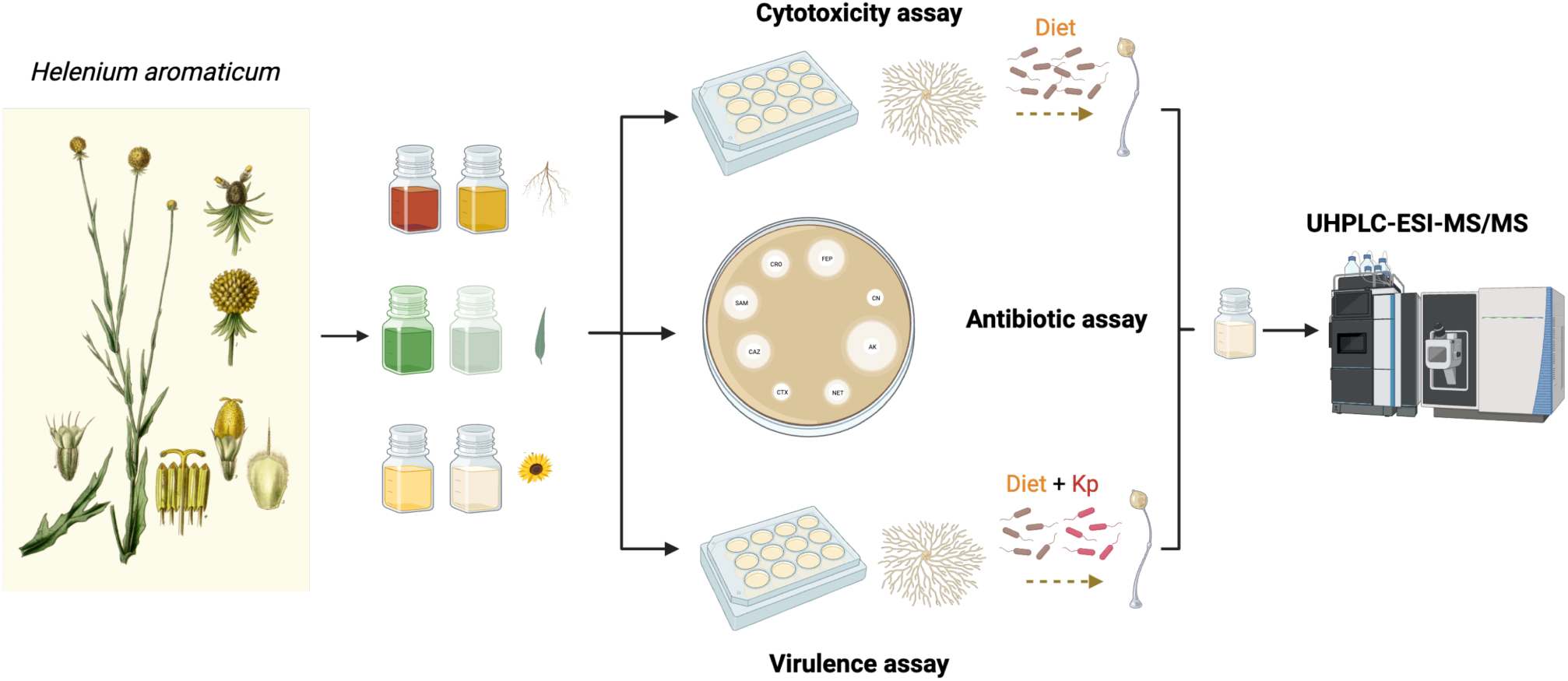

---

Antibiotic resistance is a serious global threat to public health. The overuse and misuse of antibiotics have led to the emergence of multidrug-resistant (MDR) bacterial strains that are difficult to treat ^1^. *Klebsiella pneumoniae* is a bacterium responsible for pneumonia, urinary tract infections, and sepsis, which has become increasingly resistant to antibiotics, including carbapenems, the last line of defense against MDR *K. pneumoniae* infections. The emergence of carbapenem-resistant *K. pneumoniae* strains has been a major challenge in clinical settings, as few effective treatment options are available. Antivirulence therapy is an emerging strategy for combating antibiotic resistance ^2^.

Contrary to traditional antibiotics that target essential cellular functions, antimicrobial agents focus on bacterial virulence factors critical for infection but not for bacterial growth and development. Antivirulence agents provide multiple benefits compared to conventional antibiotics; these include reducing the selection pressure leading to resistance and minimizing side effects often linked with broad-spectrum antibiotics^2^. This perspective on novel antimicrobial agents makes exploring natural resources even more pertinent.

In pharmacognosy, identifying and characterizing bioactive compounds from natural sources such as plants, fungi, and marine organisms are vital for drug discovery and therapeutic applications. Plant resources have been explored as a potential source of antimicrobial agents due to their rich chemical diversity and long history of use in traditional medicine ^3,4^. Given the complex mixtures of compounds often found in these natural products, reliable assays for evaluating biological activity, toxicity, and specificity are crucial. Therefore, combining rigorous analytical methods such as high-performance liquid chromatography (HPLC) coupled with mass spectrometry with bioassays for isolating and identifying molecules with potential pharmacological efficacy is highly desirable. Phenotypic bioassays help to ensure the quality, safety, and effectiveness of herbal medicines, supplements, and pharmaceuticals derived from natural extracts. Accordingly, implementing accurate and sensitive assays is indispensable in pharmacognosy, bridging the gap between traditional ethnomedicine and modern pharmaceutical research. However, the lack of virulence-based scalable assays for plant pharmacognosy is a bottleneck for discovering novel antivirulence molecules.

*Dictyostelium discoideum*, a soil-dwelling social amoeba, has emerged as a valuable model for host-pathogen interactions and drug discovery ^5^. *D. discoideum* is amenable to high-throughput screening of natural product extracts and has been used as a host model system for several human pathogens, including *Mycobacterium tuberculosis* ^6^, *Legionella pneumophila* ^7^*, Pseudomonas aeruginosa* ^8–10^ *and Klebsiella pneumoniae* ^11^. Some reports have used this invertebrate model system to screen the cytotoxicity of natural extracts^12–14^; however, few have simultaneously used the social amoeba to screen potential anti-virulence drugs without cytotoxic side effects ^10^. Continuing with this effort, in this study, we established an easily scalable social developmental assay in *D. discoideum* to evaluate the antivirulence activity and cytotoxicity of extracts from the plant *Helenium aromaticum* against a virulent *K. pneumoniae* strain. The genus *Helenium* is part of the *Asteraceae* family, one of the largest botanical families with numerous species with traditional medicinal uses. It is distributed in the North and part of South America^15,16^ and particularly, *Helenium aromaticum* has shown promising therapeutic potential, particularly its sesquiterpene lactones ^15,17^.

The first objective required the preparation of extracts from various parts of the *H. aromaticum* plant for the targeted screening of bioactive molecules capable of attenuating the virulence of *K. pneumoniae* RYC 492, a strain shown to be virulent over *D. discoideum* ^11^. A total of six extracts were obtained, corresponding to the dichloromethane or methanol extracts from flowers, foliage, or roots. Second, using the social amoeba *D. discoideum* as a model organism, we assessed the cytotoxicity of all the extracts based on a previously described methodology ^11^. The assay operates on the principle that secondary metabolites from various plant extracts if cytotoxic, will impede the social development of *D. discoideum* by delaying its social cycle. Conversely, non-cytotoxic compounds will facilitate social development’s normal progression and culmination. According to this criterion, the six *H. aromaticum* extracts were evaluated. The course of social development was considered within six days under our experimental conditions, and three concentrations (25 µg/mL, 50 µg/mL, and 100 µg/mL) of the different extracts were evaluated. Usually, the amoeba completes the social cycle in 48 hours under these conditions. As shown in Figure 1, amoebae development was affected by all the dichloromethane plant extracts (flowers, foliage, and roots). By day 3, the culmination of the amoeba’s cycle was reached in the controls; however, in all dichloromethane extracts, even the initial phases of the social cycle were not observed. Based on these observations, the dichloromethane extracts probably include cytotoxic compounds, indicating that further purification would be necessary to use them as a therapeutic molecule source. On the contrary, the methanolic extracts displayed no inhibitory effects on the amoeba’s social development, indicating a reduced presence of cytotoxic compounds (Figure 2A for roots). This conclusion is also sustained for flowers and foliage extracts, as shown in Figures 1SA and 2SA, where all the tested extracts allowed a normal social development of the amoeba.

**Figure 1.**
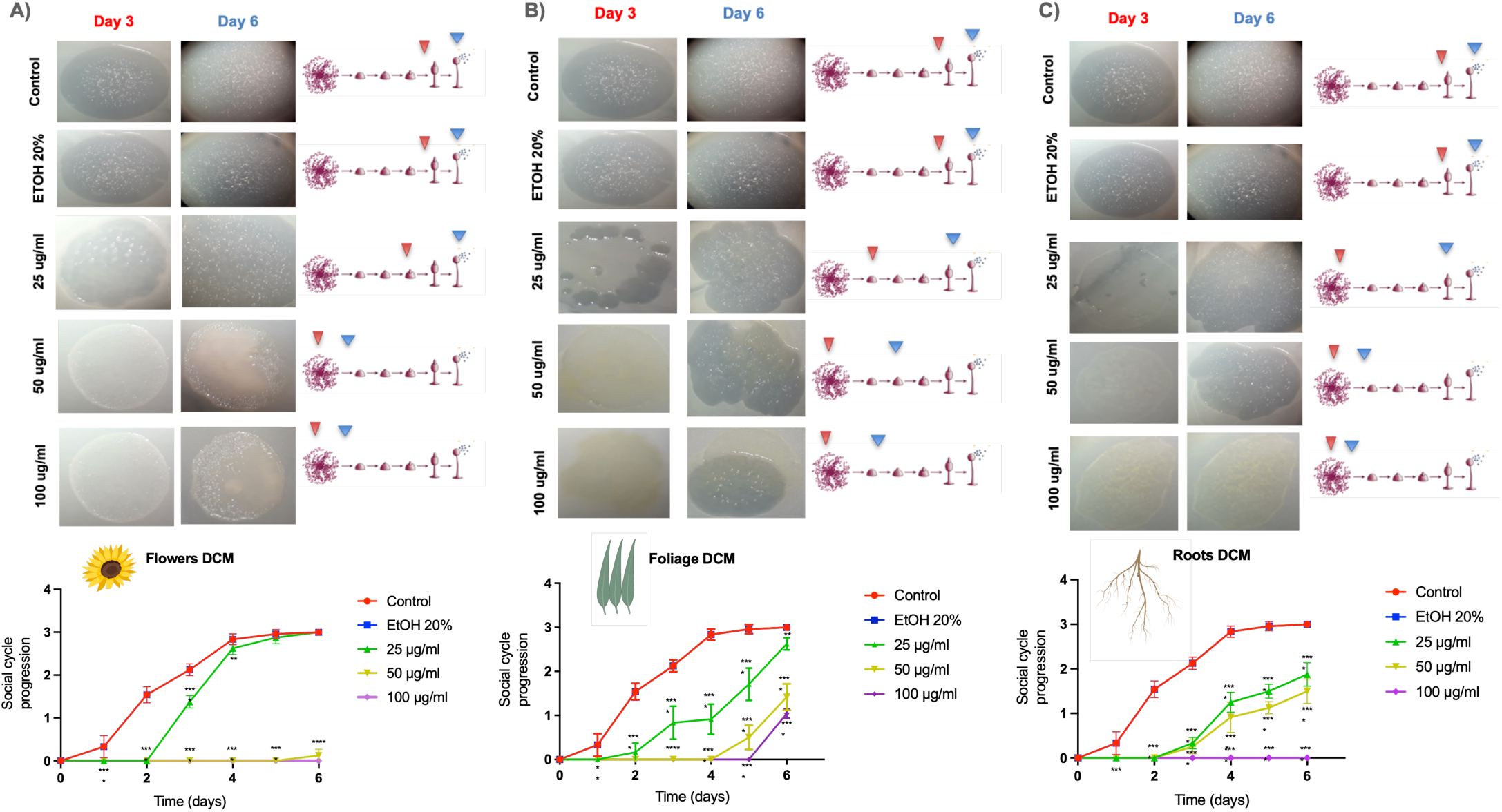
Cytotoxic evaluation of dichloromethane extracts of *H. aromaticum*. A) Flowers, B) Foliage and C) Roots. Statistical analysis was performed using a two-way ANOVA test with multiple comparisons and Dunnett’s post-test (n=9) (* = p < 0.05, ** = p < 0.005, *** = p < 0.001, ** ** = p < 0.0001). The red and blue arrowheads indicate the progression reached on day 3 and day 6, respectively. What is the control? Why EtOH 20% and not dichloromethane?

**Figure 2.**
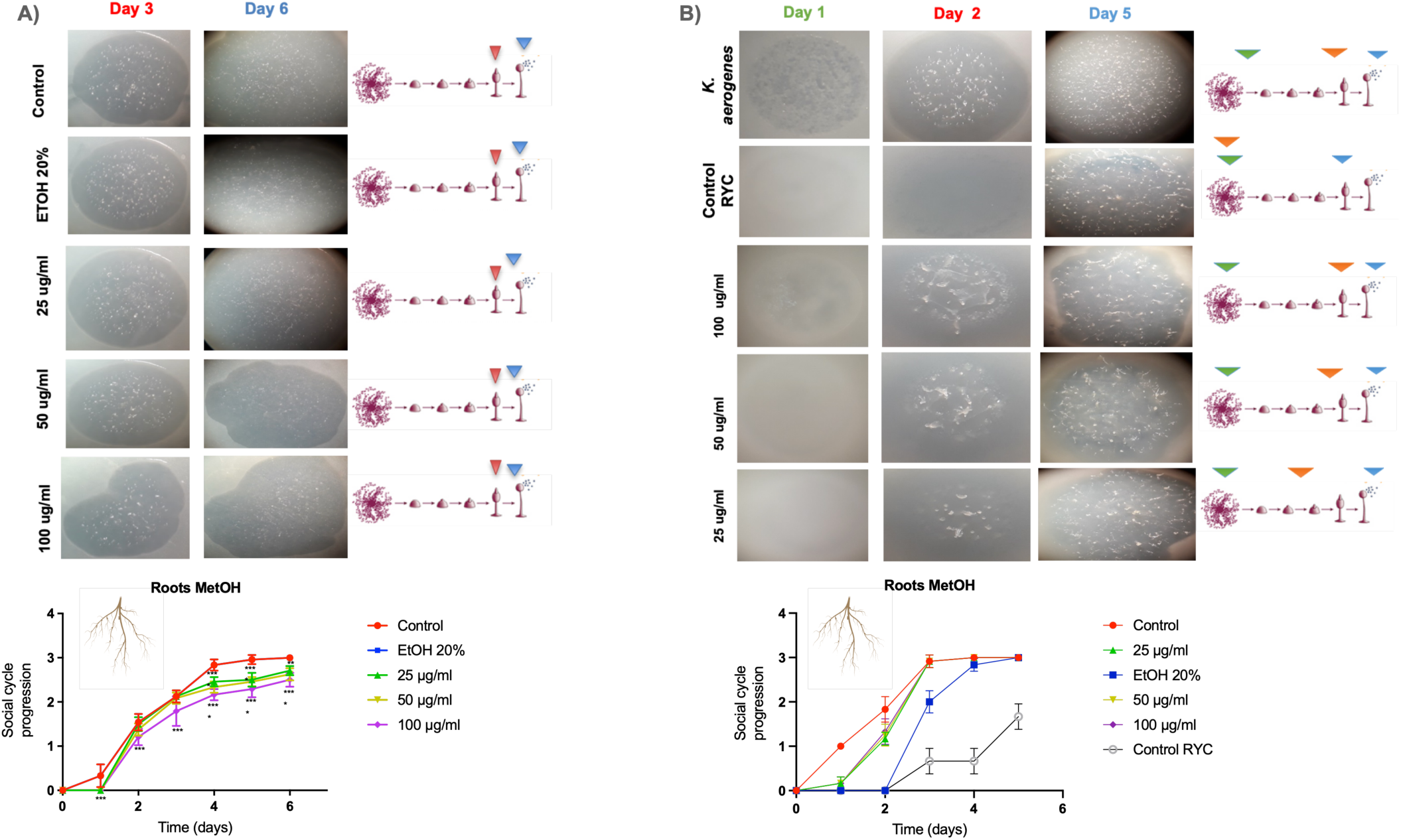
Evaluation of the cytotoxic (A) and antivirulence (B) activities of the roots methanolic extracts of *H. aromaticum*. Statistical analysis was performed using a two-way ANOVA test with multiple comparisons and Dunnett’s post-test (n=9) (* = p < 0.05, ** = p < 0.005, *** = p < 0.001, ** ** = p < 0.0001).

All methanolic extracts that showed no cytotoxic effects on the amoeba were evaluated for their antibiotic and antivirulence assets. As shown in Figure 2 and Table 1, the methanolic root extract attenuated the virulence of *K. pneumoniae* RYC492 relative to the control without compromising bacterial viability in the antibiosis assays. However, the methanolic extracts of foliage and flowers (as shown in Figures 1S and 2S and Table 1) revealed neither antibiotic nor antivirulence activity against *K. pneumoniae* RYC492. In summary, only methanolic root extract mitigated *K. pneumoniae* RYC492 virulence.

**Table 1.** Antibiosis evaluation of *H. aromaticum* extracts against the virulent *K. pneumoniae* RYC492 strain.

|  |  | Concentrations |  |  | Controls |  |  |
| --- | --- | --- | --- | --- | --- | --- | --- |
| Extracts | Bacterial species | Diameter of the inhibition zone in (mm) * |  |  |  |  |  |
| Flower dichloromethane | Klebsiella pneumoniae RYC 492 | 25 µg/ml | 50 µg/ml | 100 µg/ml | C + (KAN) resistance | C + (GEN) | C - (ETHANOL 20%) |
| Stem-leaf dichloromethane |  | - | - | +/- | +/- | +++ | - |
| Root dichloromethane |  | - | - | +/- | +/- | +++ | - |
| Flower methanolic |  | - | - | +/- | +/- | +++ | - |
| Stem-leaf methanolic |  | - | - | +/- | +/- | +++ | - |
| Root methanolic |  | - | - | +/- | +/- | +++ | - |

\* Inhibition zone, including disc diameter (6 mm) average value of repetitions n=9.
(+++) ≤ 1,8 cm, (++) ≤ 1,3 cm, (+) ≤ 0,9 cm, (+/-) ≤ 0,5 cm, (-) 0 cm, C – (negative control), C + (Kan) resistance control, C + (GEN) positive control.

Determining the chemical composition of compounds is fundamental to the pharmacognosy process, as it guides and increases the likelihood of selecting the most suitable extract for therapeutic applications. Thus, we employed high-resolution tandem mass spectrometry to identify the predominant compounds present in the extracts. A total of 63 compounds were tentatively identified (Figure 3 and Table S1), where 47 compounds were identified in the methanolic extract and 29 in the dichloromethane extract, where 13 compounds are common in both extracts. All these compounds were identified by UHPLC-Q/Orbitrap/ESI/MS/MS, which is a specific technique that identifies the compounds by their molecular weight and their characteristic fragmentations.

**Figure 3.**
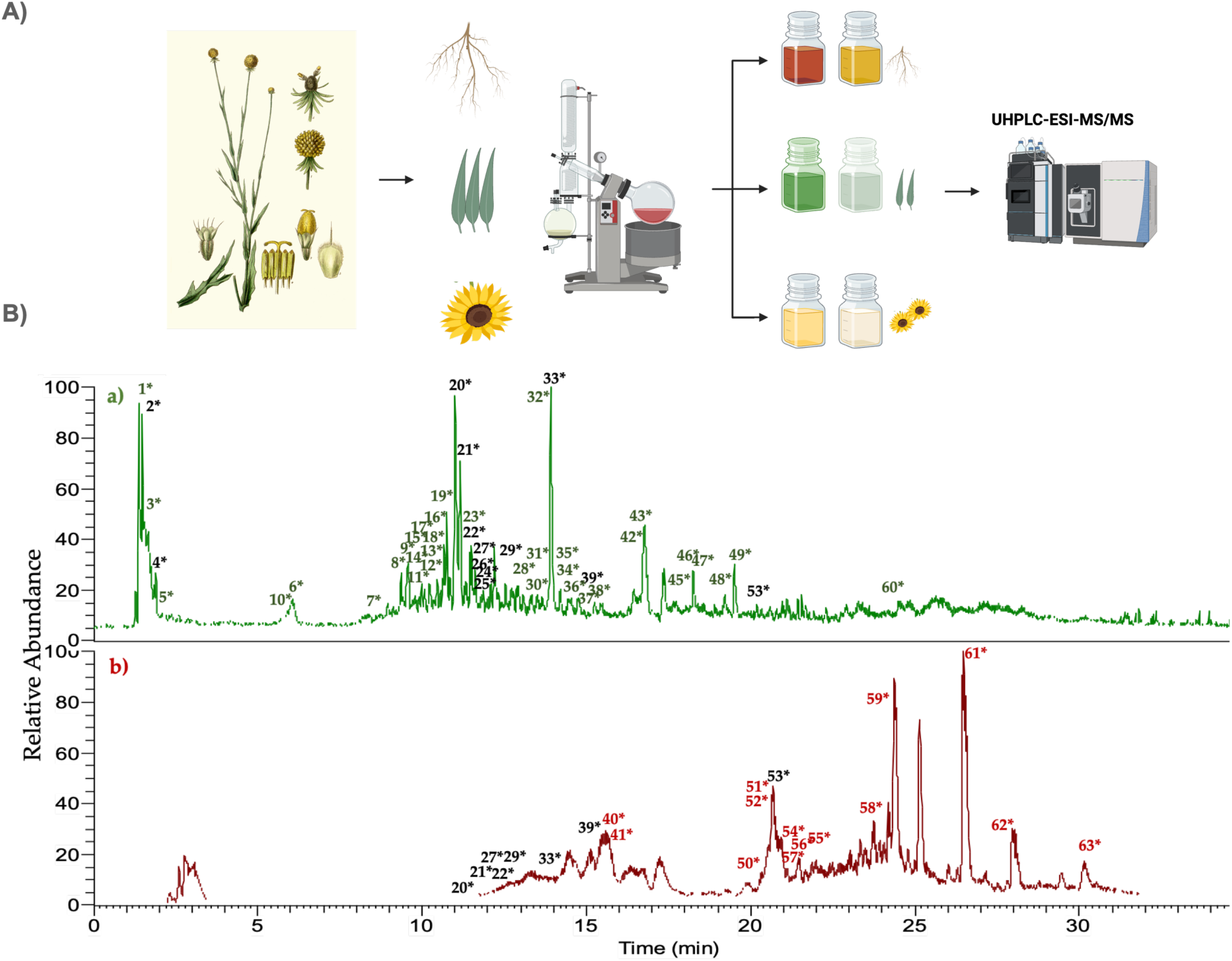
A) Overview of the extraction and identification process of *H. aromaticum* plant extracts. B) UHPLC-Q/Orbitrap/ESI/MS/MS chromatograms of the methanolic (green) and dichloromethane (red) extracts from the root of *H. aromaticum*.

Regarding phenolic compounds, 9 were identified in the methanolic extract and one in the dichloromethane extract. These compounds include dihydroxybenzoic pentoside and caffeic acid, chlorogenic acid. Fourteen flavonoids were identified, encompassing glycosylated derivatives of kaempferol, among others. Eleven terpenes were identified, primarily sesquiterpenic lactones, including various helenalin derivatives. Peaks corresponding to glycosylated terpenes, such as 10-hydroxyloganin and loganin, were also found. Other compounds, such as organic acids (succinic acid) and carbohydrates (aldopentose), also include six lipids, such as azelaic acid and pinellic acid. Sixteen compounds remained unidentified.

In summary, of the 63 compounds identified, the predominant groups were phenolic compounds, including terpenoids and flavonoids, which are likely associated with observed biological activity. For instance, sesquiterpene lactones, such as helenalin and its derivatives, might be responsible for the observed cytotoxicity, as previously reported ^18,19^. On the other hand, some flavonoids and terpenes can be accountable for attenuated virulence, as previously reported for *Klebsiella pneumoniae* ^20^.

Combining the phenotypic assays, which included antibiotic, cytotoxic, and antivirulence evaluations, with chemical analysis can be crucial to discern both the antivirulence and cytotoxic components within the plant extracts. However, further experiments with purified chemical candidates should be performed to conclusively ascertain the identity of the molecules responsible for these observed activities.

## CONCLUSIONS

Our study developed a platform to identify secondary metabolites or natural plant extracts from *H. aromaticum* that attenuated *K. pneumoniae* virulence without affecting *D. discoideum* viability. The amoeba-assisted pharmacognosy of plant resources valuation revealed that only methanolic extract from *H. aromaticum* roots fulfilled these criteria. Our approach offers a sustainable method to discover novel antivirulence agents, potentially leading to more targeted therapies with reduced resistance risks compared to traditional antibiotics.

## EXPERIMENTAL SECTION

### Microbial strains and growth conditions

Various bacterial and amoebae strains were used in this work. The *Klebsiella pneumoniae* RYC492 strain, recognized for producing the antibacterial peptide microcin E492 and salmochelin siderophores and exhibiting resistance to both kanamycin and ampicillin, was used for virulence assays. This strain was shown to be virulent over *D. discoideum*^21^. Routinely, *Klebsiella aerogenes* DBS0305928 strain was used to support the growth of *D. discoideum* ^22^. For our amoebae experiments, we used the *D. discoideum* axenic strain AX4 (DBS0302402), which emerged as a spontaneous mutation from its parental strain AX3 and was obtained from DictyBase Stock Center ^23^.

### Social development assays for cytotoxicity and *K. pneumoniae* virulence evaluations in *D. discoideum*

For the social development assays, SM agar plates embedded with a lawn of *K. aerogenes* (pre-incubated at 37°C for 18 h) were employed ^11^. Cytotoxic evaluations of the six plant extracts were conducted at 25, 50 and 100 μg/mL concentrations in 20% ethanol. Finally, the same 20% ethanol solution was used as a control growth. Briefly, 5 μL of an axenic *D. discoideum* cell suspension from selected subcultures (S3 to S5), at an exponential phase density of 2×10^6^ cells/mL, was used, totaling an inoculation of 10,000 amoebae cells. Plates were incubated at 23° C, and social development was monitored for 6 days, tracking aggregation, elevation, and culmination phases ^24^. Scoring ranged from “1” for aggregated amoebae forming a phagocytosis plaque to “3” for fully formed fruiting bodies, with transitions scored as half-values. Observations were made every 24 hours over 6 days, and the data were plotted and analyzed using GraphPad Prism.

### *H. aromaticum* collection, extracts preparation and compound identification

The plant species were collected from San Carlos de Apoquindo, Las Condes, Metropolitan Region, Chile, from January to February 2018 at latitude (S): −33.40693, longitude (W): −70.50047. The species was identified by Gloria Rojas Villegas, the Head of the Botanical Area and Herbarium at the National Museum of Natural History of Chile. The specimens were processed, deposited at the museum, and identified as *H. aromaticum* (Hook.) L.H. Bailey with a voucher number 169131.

The plant samples were oven-dried at 37°C, ground, and macerated with dichloromethane and methanol at room temperature for 72 h, repeating the process thrice with direct sonication for 1 minute and centrifugation at 10,000 xg. The extracted product was filtered and concentrated using a rotary evaporator at 40°C, then stored at −20°C. For compound identification in *H. aromaticum* extracts with either potential antivirulence or cytotoxic activity, an Ultra-High Pressure Liquid Chromatograph coupled with a diode array detector and high-resolution mass spectrometer (UHPLC-ESI-MS/MS) was employed. Sample preparation involved macerating ∼3 g of the species with methanol (three times, 30 ml each, for 3 days/extraction) to yield 10 mg of the extract.

## ASSOCIATED CONTENT

### Supporting Information

The following files are available free of charge.

Supplementary figures S1 and S2 (PDF)

Supplementary Table 2 (PDF)

## AUTHOR INFORMATION

### Corresponding Author

* Francisco P. Chávez, Phone: +56229787185

### Author Contributions

The manuscript was written through the contributions of all authors. FC and AM conceived the study. FC, AM, MH, MV and CA designed the experiments. MH, DR, MV and FC analyzed and interpreted the data. MH conducted the experiments. MH and FC wrote the manuscript. FC, MH, AM, and CA critically reviewed the manuscript. All the authors approved the final version of the manuscript.

## ACKNOWLEDGMENT

This work was funded by Fondecyt Grants 1211852 (FCH), 1221193 (AM) and 1220075 (CA). MH received the scholarship “Presidente de la República” from the Peruvian Government. We thank Nicole Molina for her excellent technician skills in preparing *D. discoideum* cultures.

## ABBREVIATIONS

DCM: dichloromethane
UHPLC: Ultra-High Pressure Liquid Chromatograph
ESI-MS/MS: Electrospray Ionization Mass Spectrometry
MDR: Multi-Drug resistant
EtOH: Ethanol.

