## Supplementary material for "*Dictyostelium discoideum-*assisted pharmacognosy of plant resources for discovering antivirulence molecules targeting *Klebsiella pneumoniae*"

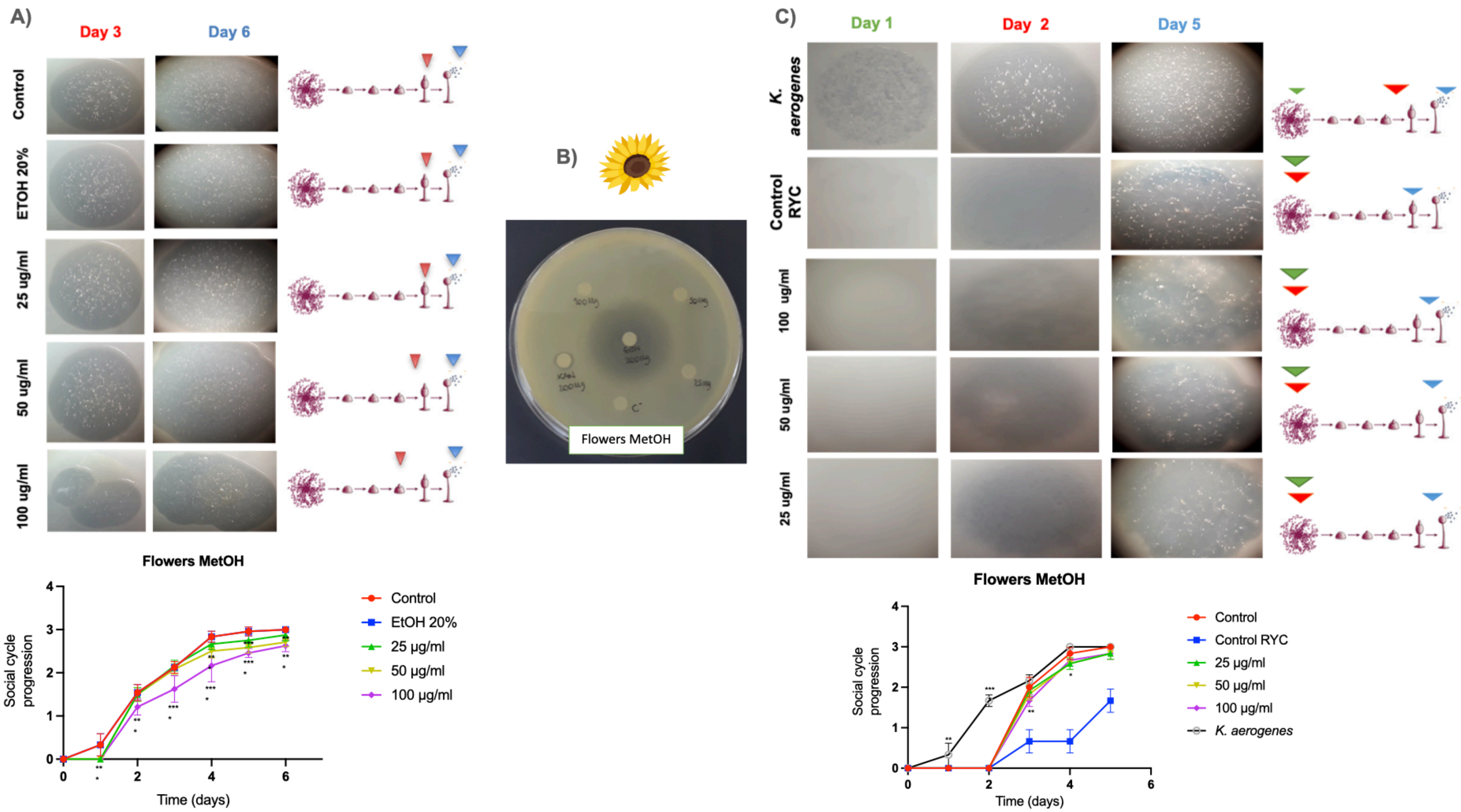


**Figure S1**. Evaluation of the cytotoxic (A) and antivirulence (B) activities of the flower methanolic extracts of *H. aromaticum*. Statistical analysis was performed using a two-way ANOVA test with multiple comparisons and Dunnett's post-test (n=9) (* = $p < 0.05$, ** = $p < 0.005$, *** = $p < 0.001$, ** ** = $p < 0.0001$).

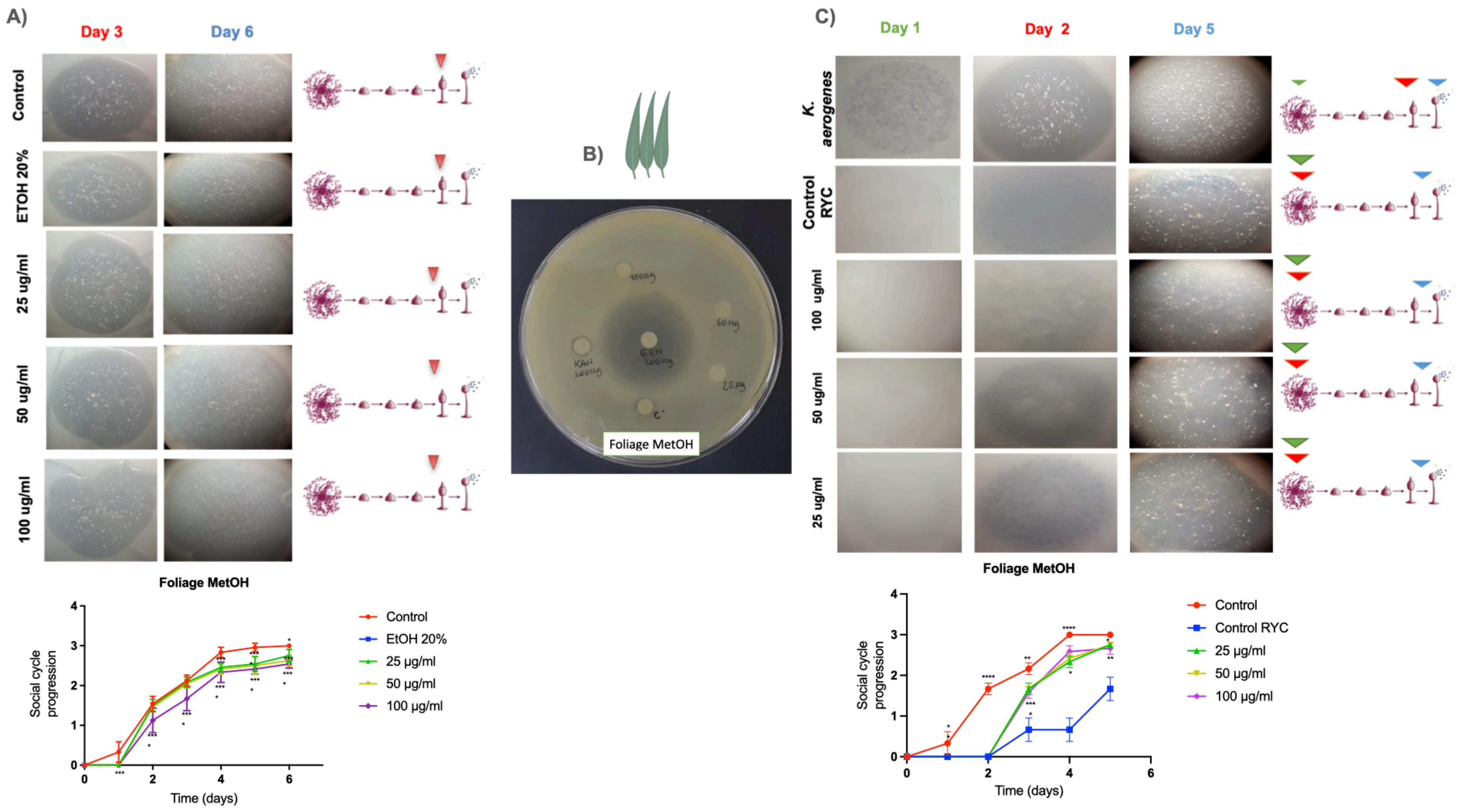


**Figure S2**. Evaluation of the cytotoxic (A) and antivirulence (B) activities of the foliage methanolic extracts of *H. aromaticum*. Statistical analysis was performed using a two-way ANOVA test with multiple comparisons and Dunnett's post-test (n=9) (* = p < 0.05, ** = p < 0.005, *** = p < 0.001, ** ** = p < 0.0001).

**Table S1. Compound comparison of the methanolic and dichloromethane extract of *H. aromaticum* root by UHPLC Q/Orbitrap/ESI/MS/MS.**

| Peak | Tentative identification | λmax (nm) | $[M-H]^-$ | RT (min) | Theoretical Mass (*m/z*) | Measured Mass (*m/z*) | Accuracy (ppm) | $MS^2$ ions (ppm) | Extract | Metabolite type |
|---|---|---|---|---|---|---|---|---|---|---|
| 1 | Vaccihein A | 226,266 | $C_{18}H_{17}O_9$ | 1.38 | 377.0873 | 377.0865 | 2.1 | 347.0757; 289.0697; 125.0237; 195.0509 | Met | Phenolic compound |
| 2 | Aldopentose | 232 | $C_5H_9O_5$ | 1.48 | 149.0450 | 149.0451 | 0.7 | - | Met; DCM | Carbohydrate |
| 3 | Syringaldehyde syringate | 271,324 | $C_{18}H_{17}O_9$ | 1.56 | 377.0873 | 377.0866 | 1.9 | 349.0914 | Met | Phenolic compound |
| 4 | Citric acid or isocitric | 228 | $C_6H_7O_7$ | 1.89 | 191.0192 | 191.0197 | 2.6 | 111.0080 | Met; DCM | Organic acid |
| 5 | Succinic acid | 226 | $C_4H_5O_4$ | 2.10 | 117.0188 | 117.0188 | 0.0 | - | Met | Organic acid |
| 6 | 10-Hydroxyloganin | 270 | $C_{17}H_{25}O_{11}$ | 6.06 | 405.1397 | 405.1413 | 3.9 | 179.0557; 163.0589 | Met | Terpene glycoside |
| 7 | Dihydroxybenzoic acid pentoside | 228,276 | $C_{12}H_{13}O_8$ | 8.75 | 285.0610 | 285.0625 | 5.3 | 108.0209 153.0189 | Met | Phenolic compound |
| 8 | Chlorogenic acid | 241,325 | $C_{16}H_{17}O_9$ | 9.36 | 353.0873 | 353.0888 | 4.2 | 191.0561 | Met | Phenolic compound |
| 9 | Kaempferol-3-O-hexoside-pentoside | 232,271,326 | $C_{27}H_{29}O_{15}$ | 9.55 | 593.1506 | 593.1526 | 3.4 | 431.0988 161.0239 | Met | Flavonoid |
| 10 | Loganin | 271,324 | $C_{17}H_{25}O_{10}$ | 9.59 | 389.1448 | 389.1464 | 4.1 | 359.1358; 343.1407; 227.0930 | Met | Terpene glycoside |
| 11 | Esculetin | 269,319 | $C_9H_5O_4$ | 9.74 | 177.0188 | 177.0193 | 2.8 | 133.0288 121.0289 | Met | Coumarin |

| | | | | | | | | | | |
|---|---|---|---|---|---|---|---|---|---|---|
| 12 | Caffeic acid | 245,324 | $C_9H_7O_4$ | 9.85 | 179.0344 | 179.0349 | 2.8 | 135.0446 | Met | Phenolic compound |
| 13 | Kaempferol-3-O-rutinoside | 242,271,333 | $C_{27}H_{29}O_{15}$ | 9.98 | 593.1506 | 593.1527 | 3.5 | 285.0455; 161.0242 | Met | Flavonoid |
| 14 | Unknow | 244 | $C_{20}H_{31}O_{10}$ | 10.08 | 431.1917 | 431.1933 | 4.0 | 114.0554 | Met | - |
| 15 | Unknow | 273 | $C_{20}H_{30}O_6N$ | 10.22 | 380.2073 | 380.2090 | 4.5 | 114.0553 | Met | - |
| 16 | Kaempferol-7-O-glucoside | 270,343 | $C_{21}H_{19}O_{11}$ | 10.45 | 447.0927 | 447.0944 | 3.8 | 327.0523 | Met | Flavonoid |
| 17 | Azalein | 271,329 | $C_{22}H_{21}O_{11}$ | 10.54 | 461.1084 | 461.1102 | 3.9 | 431.0992; 315.0552 | Met | Flavonoid |
| 18 | Kaempferol-7-O-glucoside isomer | 255,267,347 | $C_{21}H_{19}O_{11}$ | 10.68 | 447.0927 | 447.0943 | 3.6 | 327.0519 | Met | Flavonoid |
| 19 | Caffeoyl-xylose | 243,326 | $C_{14}H_{15}O_8$ | 10.75 | 311.0767 | 311.0781 | 4.5 | 149.0449 179.0347 | Met | Phenolic compound |
| 20 | Vitexin | 237,268,343 | $C_{21}H_{19}O_{10}$ | 11.00 | 431.0978 | 431.0994 | 3.7 | 341.0677; 311.0571 | Met; DCM | Flavonoid |
| 21 | Cytisoside | 233,271,336 | $C_{22}H_{21}O_{10}$ | 11.14 | 445.1135 | 445.1151 | 3.6 | 311.0559; 283.0619 | Met; DCM | Flavonoid |
| 22 | Cytisoside isomer | 243,269,330 | $C_{22}H_{21}O_{10}$ | 11.45 | 445.1135 | 445.1152 | 3.8 | 311.0573; 283.0612 | Met; DCM | Flavonoid |
| 23 | Gnaphaliol glucopyranoside | 257,279 | $C_{19}H_{23}O_9$ | 11.65 | 395.1342 | 395.1358 | 4.0 | - | Met | Benzofuran |
| 24 | Unknow | 270,329 | $C_{22}H_{31}O_{12}$ | 11.85 | 487.1816 | 487.1834 | 3.7 | - | Met; DCM | - |
| 25 | Unknow | 247,276,329 | $C_{14}H_{13}O_7$ | 12.20 | 293.0661 | 293.0677 | 5.4 | - | Met; DCM | - |
| 26 | Azelaic acid | - | $C_9H_{15}O_4$ | 12.27 | 187.0970 | 187.0976 | 3.2 | 125.0965 | Met; DCM | Lipid |

| | | | | | | | | | | |
|---|---|---|---|---|---|---|---|---|---|---|
| 27 | Hydroxy-dihydrohelenalin | 244,273 | $C_{15}H_{19}O_5$ | 12.54 | 279.1232 | 279.1246 | 5.0 | 263.1296; 251.1295; 235.1345 | Met; DCM | Sesquiterpene lactone |
| 28 | Ferulic acid | 274,324 | $C_{10}H_9O_4$ | 12.80 | 193.0501 | 193.0508 | 3.6 | 179.0349; 175.0398; 149.0603; | Met | Phenolic compound |
| 29 | Dihydroxy-dihydrohelenalin | 239 | $C_{15}H_{19}O_6$ | 12.89 | 295.1182 | 295.1188 | 0.3 | 251.1295 | Met; DCM | Sesquiterpene lactone |
| 30 | 6-O-Acetylglycitin | 270 | $C_{24}H_{23}O_{11}$ | 12.97 | 487.1240 | 487.1259 | 3.9 | 283.0620 | Met | Flavonoid |
| 31 | Nitrophenol | 247 | $C_6H_4O_3N$ | 13.08 | 138.0191 | 138.0192 | 1.2 | - | Met | Nitro phenolic compound |
| 32 | Kaempferol | 270,337,367 | $C_{15}H_9O_6$ | 13.82 | 285.0399 | 285.0415 | 5.6 | 151.0033; 135.0447 | Met | Flavonoid |
| 33 | Isorhamnetin | 254,272,345 | $C_{16}H_{11}O_7$ | 13.90 | 315.0505 | 315.0520 | 4.8 | 300.0286; 215.0099 | Met; DCM | Flavonoid |
| 34 | Unknow | 213 | $C_{16}H_{23}O_6$ | 14.29 | 311.1495 | 311.1502 | 2.2 | - | Met | - |
| 35 | Unknow | 211 | $C_{21}H_{17}O_2$ | 14.44 | 301.1229 | 301.1213 | 5.3 | - | Met | - |
| 36 | Unknow | 272 | $C_{25}H_{15}O_3N_2$ | 14.76 | 391.1083 | 391.1079 | 1.0 | - | Met | - |
| 37 | Unknow | 217,333 | $C_{23}H_{17}O_5$ | 15.08 | 373.1076 | 373.1060 | 4.2 | - | Met | - |
| 38 | Rhododendrin | 272 | $C_{16}H_{23}O_7$ | 15.26 | 327.1444 | 327.1459 | 4.6 | - | Met | Phenolic compound |
| 39 | 11$\alpha$, 13-Dihydrohelenalin | 270 | $C_{15}H_{19}O_4$ | 15.49 | 263.1283 | 263.1296 | 4.9 | 219.1392 | Met; DCM | Sesquiterpene lactone |
| 40 | Dihydrohelenalin glucose | 223,306 | $C_{21}H_{29}O_9$ | 15.55 | 425.1812 | 425.1816 | 0.9 | 381.1557; 263.1283 | DCM | Sesquiterpene lactone |
| 41 | Unknow | 225 | $C_{23}H_{19}O_4$ | 15.77 | 359.1283 | 359.1269 | 3.9 | | DCM | - |
| 42 | Apigenin | 267,335 | $C_{15}H_9O_5$ | 16.44 | 269.0450 | 269.0465 | 5.6 | 117.0341; 149.0240; 133.0296 | Met | Flavonoid |
| 43 | Methyl-luteolin | 273,335 | $C_{16}H_{11}O_6$ | 16.77 | 299.0556 | 299.0570 | 4.7 | 285.0362; 1510395 | Met, DCM | Flavonoid |

| | | | | | | | | | | |
|---|---|---|---|---|---|---|---|---|---|---|
| 44 | Hydroxy-dihydrohelenalin isomer | 230 | $C_{15}H_{19}O_5$ | 16.80 | 279.1232 | 279.1239 | 2.5 | 263.1296; 251.1295; 235.1345 | DCM | Sesquiterpene lactone |
| 45 | Unknow | 250 | $C_{28}H_{29}O_{13}$ | 17.69 | 573.1608 | 573.1661 | 9.2 | - | Met | - |
| 46 | Pinellic acid | 271 | $C_{18}H_{33}O_5$ | 18.24 | 329.2328 | 329.2343 | 4.6 | - | Met | Lipid |
| 47 | Acevaltrate | 270 | $C_{24}H_{31}O_{10}$ | 18.37 | 479.1917 | 479.1935 | 3.8 | - | Met | Iridoid monoterpene |
| 48 | Unknow | 271 | $C_6H_{13}O_{10}$ | 19.22 | 245.0509 | 245.0496 | 5.3 | - | Met | - |
| 49 | Unknow | 251 | $C_{30}H_{31}O_{14}$ | 19.51 | 615.1714 | 615.1768 | 8.8 | - | Met | - |
| 50 | Helenalin | 236 | $C_{15}H_{17}O_4$ | 19.89 | 261.1127 | 261.1132 | 1.9 | 233.1183; 229.0868; 219.1389; 185.0968 | DCM | Sesquiterpene lactone |
| 51 | Acetyl hydroxy-helenalin | 236 | $C_{17}H_{21}O_6$ | 20.33 | 321.1338 | 321.1346 | 2.5 | 277.1078; 261.1132; 217.1232; 173.1331 | DCM | Sesquiterpene lactone |
| 52 | Vanillin | 224,280 | $C_8H_7O_3$ | 20.52 | 151.0395 | 151.0396 | 0.6 | 133.0653 | DCM | Phenolic compound |
| 53 | Methyl isorhamnetin | 214,342 | $C_{17}H_{13}O_7$ | 20.95 | 329.0661 | 329.0668 | 2.1 | - | DCM | Flavonoid |
| 54 | Tianshic acid | - | $C_{18}H_{33}O_5$ | 21.13 | 329.2328 | 329.2336 | 2.4 | - | DCM | Lipid |
| 55 | Hydroxytetracosapentaenoic acid | - | $C_{24}H_{37}O_3$ | 21.44 | 373.2743 | 373.2750 | 1.9 | - | DCM | Lipid |
| 56 | Wallichoside | 234 | $C_{20}H_{27}O_8$ | 21.69 | 395.1706 | 395.1713 | 1.8 | - | DCM | Steroid |
| 57 | Unknow | 233,273 | $C_9H_{13}O_3$ | 21.85 | 169.0865 | 169.0868 | 1.8 | - | DCM | - |
| 58 | Unknow | 233,275 | $C_{10}H_9O_2$ | 22.00 | 161.0603 | 161.0604 | 0.6 | - | DCM | - |
| 59 | Unknow | 234,301 | $C_{30}H_{37}O_9$ | 23.06 | 541.2438 | 541.2435 | 0.5 | - | DCM | - |
| 60 | Tetrahydrohelenalin | 272 | $C_{15}H_{21}O_4$ | 24.55 | 265.1440 | 265.1459 | 7.2 | - | Met | Sesquiterpene lactone |
| 61 | Hydroxyoctadecadienoic acid | - | $C_{18}H_{31}O_3$ | 26.46 | 295.2273 | 295.2278 | 1.7 | - | DCM | Lipid |
| 62 | Unknow | 249 | $C_{30}H_{31}O_{14}$ | 27.15 | 615.1714 | 615.1758 | 7.1 | - | DCM | - |

| | | | | | | | | | | |
|---|---|---|---|---|---|---|---|---|---|---|
| 63 | Hydroxytetracosapentaenoic acid isomer | - | $C_{24}H_{37}O_3$ | 30.14 | 373.2743 | 373.2747 | 1.1 | - | DCM | Lipid |
